# ACE1 Does Not Influence Cerebral Aβ Degradation or Amyloid Plaque Accumulation in 5XFAD Mice

**DOI:** 10.1101/2024.09.27.615540

**Authors:** Sohee Jeon, Alia O. Alia, Jelena Popovic, Robert J. Vassar, Leah K. Cuddy

## Abstract

Alzheimer’s disease (AD) is the most common form of dementia, and multiple lines of evidence support the relevance of Aβ deposition and amyloid plaque accumulation in the neurotoxicity and cognitive decline in AD. Rare mutations in angiotensin-converting-enzyme-1 (ACE1) have been highly associated with late onset AD patients; however, the mechanism for ACE1 mutation in AD pathogenesis is unknown. Given the relevance of ACE1 with AD and the strong association of Aβ to AD pathogenesis, we investigated whether ACE1 degrades Aβ and affects amyloid burden in 5XFAD mice *in vivo*. To investigate this, we analyzed 6-month-old 5XFAD mice with ACE1 loss of function. ACE1 loss of function was mediated either by crossing 5XFAD mice to ACE1 conditional knockout mice or administering 5XFAD mice with the ACE1 inhibitor enalapril. Our analyses revealed that ACE1 loss of function through both genetic and pharmacological methods does not affect amyloid plaque load and neuroinflammation in the hippocampus and cortex of 5XFAD mice.

## Introduction

Alzheimer’s disease (AD) is a neurodegenerative disorder that affects around 6.9 million Americans ages 65 and older in the United States^1^. AD initially manifests as a decline in memory and cognition, and eventually, leads to death^1,2^. Distinct from other forms of dementia, the pathological hallmarks of AD consist of amyloid plaques and neurofibrillary tangles^3^. Specifically, amyloid plaques consist primarily of amyloid beta peptides (Aβ), particularly the neurotoxic Aβ42, which initially accumulates in neurons, and subsequently aggregates extracellularly in cerebral cortex, subiculum, hippocampus, and other regions of the brain as the disease progresses^3–5^. Multiple lines of evidence support the relevance of Aβ deposition and amyloid plaque accumulation in the neurotoxicity and cognitive decline in AD^4,6,7^. Furthermore, amyloid plaque accumulation has been linked to mutations in APP, PSEN1, and PSEN2; these mutations lead to familial autosomal dominant or early onset AD (EOAD) which comprise less than 5% of AD cases^7,8^. Symptomatically indistinguishable from EOAD, late onset AD (LOAD) comprises for most AD cases and associates with mutations such as APOE and TREM2^3^.

Genome-wide association studies (GWAS) on AD patients revealed additional genetic mutations highly associated with LOAD, including those in ACE1^9,10^. ACE1 encodes for angiotensin-converting-enyme-1, an enzyme that cleaves angiotensin I (AngI) to angiotensin II (AngII) for activation of various receptors, including AngII receptor type 1 (AT1R) and AngII receptor type 2 (AT2R), in the renin-angiotensin-system (RAS)^11,12^. While the peripheral RAS has been well known for its role in blood pressure control, the central RAS has been linked to neurocognition^12–15^. Furthermore, most analysis of postmortem brain, CSF, and plasma from AD patients reveal alterations in ACE1 level and/or activity, further suggesting a connection between AD and ACE1^9,10,16,17^. In particular, a study identified the ACE1 R1279Q mutation in AD families through whole genome sequencing, and its cognate mutation in knock-in (KI) mice caused age-dependent brain atrophy and hippocampal neurodegeneration^10^. This study further examined 5XFAD mice with the ACE1 KI mutation and reported that, while Aβ accelerated neurodegeneration, Aβ42 levels remained unchanged in these mice^10^. Given the relevance of ACE1 with AD and the strong association of Aβ to AD pathogenesis, multiples studies sought to understand whether ACE1 affects Aβ deposition and amyloid accumulation.

Past studies investigating the role of ACE1 in AD suggest that ACE1 cleaves and degrades Aβ^18,19^. Although few *in vivo* studies support the hypothesis that ACE1 contributes to Aβ degradation, *in vitro* studies have reported that the N-domain of ACE1 specifically cleaves Aβ^20–24^. While *in vitro* studies and some *in vivo* studies imply ACE1’s role in Aβ cleavage and degradation^19,25–27^, most clinical and *in vivo* data suggest otherwise^17,28^. For example, several clinical studies suggest that administration of centrally acting ACE1 inhibitors to AD patients delays or prevents cognitive decline, improves cognition, and reduces the progression of disease neuropathology; thus, inhibiting ACE1 may have clinical benefits^29–36^. Adding to this conundrum, several *in vivo* and clinical studies suggest that ACE1 does not affect Aβ levels in the brain and further report no significant difference in AD incidence with ACE1 inhibitors^37–44^. Altogether, these findings suggest conflicting results regarding the role of ACE1 in Aβ degradation. However, discrepancies may arise due to factors such as study design, the limited ability of ACE1 inhibitors to cross the blood-brain barrier and their effects on non-neuronal cells, despite ACE1 being predominantly expressed in neurons.

Here, we investigated whether ACE1 degrades Aβ and affects amyloid burden in the AD brain *in vivo*. We developed a novel mouse model where ACE1 is specifically knocked out in forebrain neurons of the 5XFAD mouse model of amyloidosis. Investigating whether ACE1 catalyzes Aβ is important to not only understand the mechanism of ACE1 in AD pathogenesis, but also to examine the neurocognitive effect of ACE inhibitors as a treatment for hypertensive patients. For this, we examined amyloid burden in 5XFAD amyloid transgenic mice with ACE1 inhibition, and we hypothesized that ACE1 does not impact Aβ levels *in vivo* based on previous transgenic AD mouse studies.

## Methods

### Mouse Brain Extraction

Mice were measured for their body weight and were subsequently euthanized by intraperitoneal injection of ketamine (100 mg/kg) and xylazine (15 mg/kg). Then, mice were perfused with the Perfusion solution (20 µg/ml phenylmethyl sulfonyl fluoride, 0.1mM dithiothreitol, 0.5 µg/ml leupeptin, and 20µM sodium orthovanadate in 1x PBS). This was immediately followed by brain removal and brain weight measurement. Brains were divided in half along the sagittal plane for hemibrains which were immediately preserved for downstream experiments. Hemibrains from perfused mice were fixed in 10% formalin and preserved in 30% sucrose 1x PBS solution at 4°C. For experiments requiring analysis of specific brain regions, hemibrains were dissected on ice for the hippocampus, cortex and cerebellum and flash frozen in liquid nitrogen for storage at −80°C until use.

### Animals

All animal work was done in accordance with Northwestern University Institutional Animal Care and Use Committee (IACUC) approval. Mice were fed a standard rodent chow diet and water ad libitum, and housed with a standard 12-hour light/dark cycle. 5XFAD mice on a C57BL6 background (Jackson Laboratories) were maintained by crossing transgene positive hemizygous male C57BL6 mice with transgene negative female C57BL6 mice (Jackson Laboratories). To generate ACE1 conditional knockout mice, the targeting strategy was based on NCBI transcript NM_207624.5 which corresponds to Ensembl transcript ENSMUST00000001963 (Ace-001). The positive selection marker (Puromycin resistance, PuroR) was flanked by FRT sites and was inserted into intron 13. The targeting vector was generated using BAC clones from the C57BL/6J RPCI-23 BAC library and transfected into the Taconic Biosciences C57BL/6N Tac ES cell line. Homologous recombinant clones were isolated using positive (PuroR) and negative (Thymidine kinase, Tk) selections. The conditional KO allele was obtained after in vivo Flp-mediated removal of the selection marker. Deletion of exons 14 and 15 resulted in the loss of function of the *Ace* gene by deleting part of the Extra Cellular Domain and by generating a frameshift from exon 13 to exons 16-18, a premature stop codon is in exon 16. The remaining recombination site is located in a non-conserved region of the genome. Cre driver mice that express cre-recombinase under the control of the CamKIIα promoter (Jackson Laboratories) were used to generate 5XFAD; ACE1 cKO mice lacking ACE1 expression specifically in excitatory forebrain neurons.

Transgene positive and negative ACE1^FL/+^ mice were crossed to ACE1^FL/+^ heterozygous CamKIIα-iCre mice to generate ACE1^FL/FL^ iCre mice, as well as ACE1^FL/FL^, ACE1^+/+^iCre and ACE1^+/+^ littermates, which served as controls for the study.

#### Drug Treatment

Enalapril (Sigma-Aldrich) was prepared at 3 g/L in drinking water, providing a daily dose of approximately 300 mg/kg—about twice the maximum dose for hypertensive patients after body surface area conversion. Treatment with enalapril or vehicle started in 5XFAD mice at 2 months old, when amyloid plaques begin forming in the 5XFAD model, and continued until 6 months. This duration was chosen to observe any preventative effect on plaque formation. Fresh drug solution was prepared, and water was changed every 48 hours.

### Enalaprilat Concentration Measurement

The development of enalaprilat measurement methods and the corresponding analysis were executed by Pharmaron (Exton) Lab Services LLC as follows: Each brain sample was weighed, combined with 9 volumes of 20% methanol in water, and thoroughly homogenized. Blank mouse brain (BioIVT, Lot# MSE495285, 495287, 495289, 495293, 495301) was weighed and combined with 9 volumes of 20% methanol in water, homogenized, and used as the matrix for preparation of calibration standards. Aliquots of brain homogenate (50 μL) were combined with 100 μL of acetonitrile in a 96-well plate. Calibration standards were prepared in homogenized brain and immediately combined with 9 volumes of acetonitrile. Calibration standards ranged from 1 μg/mL to 0.3 ng/mL. After thorough mixing, the plate was centrifuged for 10 minutes at 2242 x g. In a separate plate, an aliquot of supernatant (100 μL) was removed, diluted with one volume of water containing internal standard (0.02 μM enalaprilat-d_5_), and analyzed by LC-MS/MS.

### Immunoblotting Assay

#### Protein extraction

Tissues were weighed and kept on dry ice for sequential extraction into PBS and RIPA fractions. For the PBS extraction, tissues were manually homogenized with a hand-held homogenizer in PBS extraction buffer [Halt phosphatase inhibitor (#78420, Thermo Fisher Scientific) and Protease inhibitor cocktail III (#535140, EMD Millipore) in 1x PBS buffer] at 1:10 (w/v). Homogenates were centrifuged at 14,000 RPM for 30 minutes at 4°C, and the supernatant was removed as PBS-soluble fraction for storage at −80°C until use. The pellet was further extracted by the addition of RIPA extraction buffer [Halt phosphatase inhibitor, Protease inhibitor cocktail III, 50mM tris, 0.15M NaCl, 1% IGEPAL, 1mM EDTA, 1mM EGTA, 0.1% SDS and 0,5% sodium deoxylate in nanopure water, adjusted to pH 8.0] at 1:10 (w/v) and incubation on ice for 45 minutes. This was followed by sonication (Misonix XL-2000) at setting ‘5’ for 20 seconds on an ice-slurry and centrifugation at 14,000 RPM for 30 minutes at 4°C. The supernatant was removed as the RIPA-soluble fraction and stored at −80°C until use.

#### Western blot

Sample protein concentrations were measured using bicinchoninic acid (BCA) assay (#23225, Thermo Fisher Scientific) followed by preparation of western blot samples by adding 4X NuPAGE LDS sample buffer (#NP0008, Thermo Fisher Scientific) with 4% β-Mercaptoethanol (#M6250, Millipore Sigma) to extracted proteins and heating at 95°C for 10 minutes. Samples containing equal amounts of protein were loaded into NuPAGE 4-12% midi Bis-tris gels (#WG1403BOX, Thermo Fisher Scientific) for running in 1x MOPS running buffer [50mM MOPS (#PHG0007, Millipore Sigma), 50mM Tris Base (#DST60040-10000, Dot Scientific), 1mM EDTA (#50-841-667, Teknova) and 0.1% SDS (#50-751-6948, Quality Scientific) in nanopure water] by using the Criterion Cell electrophoresis chamber (Bio-rad). Then, proteins were transferred to nitrocellulose membrane in 5x transfer buffer (Bio-rad) for 45 minutes at 1.1A and 25V using the Trans-Blot Turbo Transfer system (Bio-rad). The membrane was washed 3 times in Wash buffer (0.1% Tween-20 in 1x PBS) for 5 minutes per wash, and was blocked in SuperBlock blocking buffer (#37517, Thermo Fisher Scientific) at room temperature for 1 hour. Subsequently, the membrane was incubated with Primary antibody solution (Primary antibody and 10% SuperBlock blocking buffer in Wash buffer) overnight at 4°C. Next day, the membrane was washed 3 times in Wash buffer for 5 minutes per wash and was incubated with Secondary antibody solution (Secondary antibody and 10% SuperBlock blocking buffer in Wash buffer) at room temperature for 1 hour. The membrane was then washed 3 times in Wash buffer for 5 minutes at room temperature and developed with SuperSignal West Pico PLUS Chemiluminescent Substrate (#34580, Thermo Fisher Scientific) or SuperSignal West Femto Maximum Sensitivity Substrate (#34096, Thermo Fisher Scientific). Finally, membranes were imaged on a Bio-rad Chemidoc MP Imaging System for quantification analysis with the ImageLab Version 6.1 software.

### Enzyme-linked Immunosorbent Assay (ELISA)

Angiotensin II EIA Kit (#RAB0010, Sigma-Aldrich) was performed to detect ang II peptide levels. The assay was performed in accordance with the manufacturer provided protocol. In brief, 100µl of diluted AngII antibody was added to each well of the precoated plate and was incubated overnight at 4°C with gentle shaking (1-2 cycles/second). Plates were then wash 4 times with 200µl of wash buffer per well. Next, 100µl of the protein samples, standards, positive control, or negative control was added to appropriate wells with a blank well. Each protein sample was prepared by combining 125µl of undiluted PBS-soluble protein sample (refer to the following methods section: ‘Immunoblotting assay’ – ‘Protein extraction’). with 125µl of biotinylated AngII Peptide at 40 pg/ml. Positive control was prepared by mixing 100µl of AngII Positive Control Sample, 100µl of biotinylated AngII Peptide at 40 pg/ml, and 4µl of 10-fold diluted biotinylated AngII Peptide. Negative control was prepared by combining 125µl of 1x PBS with 125µl of biotinylated AngII Peptide at 40 pg/ml. Standards were prepared at 1000, 100, 10, 1, 0.1, and 0 pg/ml by diluted EIA AngII Peptide standard with biotinylated AngII Peptide Working Stock at 20 pg/ml. Following this, the wells were covered with a film and incubated overnight at 4°C with gentle shaking (1-2 cycles/second). Next day, plates were washed 4 times with 200µl of wash buffer per well, followed by addition of 100µl of HRP-Streptavidin solution to each well for 45 minutes incubation with gentle shaking at room temperature. Plates were washed 4 times again with 200µl of wash buffer per well, and then, 100µl of TMB One-Step Substrate Reagent was added to each well for 30 minutes incubation at room temperature with gentle shaking. Lastly, 50µl of Stop Solution was added to each well and the plates were immediately measured for absorbance at 450nm.

### Immunohistochemistry

#### Sectioning

Hemibrains were frozen and sectioned at 30-microns thickness along the coronal or sagittal plane using a freezing-sliding microtome. These sections were serially harvested into a 12-well plate with cryoprotective solution (1x PBS, 30% sucrose, and 30% ethylene glycol) and preserved for long-term storage at −20°C.

#### Staining for Immunofluorescence

Brain sections from specific Bregma position were selected for each immunofluorescence analysis: NeuN, Aβ42, Iba1 and GFAP analysis (Bregma coordinates of approximately −1.70 to −3.52mm) and brain volume analysis (Bregma coordinate of approximately −0.94 to −3.40mm). Selected free-floating sections were placed into individual wells in a non-treated 24-well plate and were washed three times (5 minutes incubation per wash) in 1x tris-buffered saline (TBS) at room temperature with gentle shaking on an orbital shaker. Sections were then incubated in Glycine solution (16mM glycine and 0.25% Triton X-100 in 1x TBS) for 1 hour at room temperature with gentle shaking and were washed three times (5 minutes incubation per wash) in 1x TBS with gentle shaking. Next, sections were incubated in Blocking solution (0.25% Triton X-100, 5% donkey serum in 1x TBS) for either 2 hours at room temperature or overnight at 4°C with gentle shaking and were washed two times (10 minutes incubation per wash) in BSA solution (1% BSA and 0.25% Triton X-100 in 1x TBS). Sections were incubated in primary antibodies solution (primary antibodies, 1% BSA and 0.25% Triton X-100 in 1x TBS) overnight at 4°C with gentle shaking and were then washed three times (10 minutes incubation per wash) in BSA solution.

Subsequently, sections were incubated in secondary antibodies solution (secondary antibody, 1% BSA and 0.25% Triton X-100 in 1x TBS) for 2 hours at room temperature with gentle shaking in the dark and were then washed three times (15 minutes incubation per wash) in 1x TBS. All sections were mounted onto slides (#MCOMM/W/90, StatLab) using ProLong Gold (#P36934, Thermo Fisher Scientific) with coverslips (#63791-01, Electron Microscopy Sciences) and dried overnight at room temperature.

#### Imaging and Quantification for Volumetric Analysis

Mounted anti-NeuN stained sections were imaged on the Ti2 wide-field microscope (Northwestern University Center for Advanced Microscopy and Nikon Imaging Centre). Sequential sections that were 360µm apart within Bregma coordinate of approximately −0.94 to −3.40mm were selected for volumetric analysis. ImageJ software was used to trace and measure the area of interest (hippocampus, cortex, or cerebellum) on the section images. Finally, volume was calculated using the formula: volume = (sum of area) x 0.36mm.

#### Imaging and Quantification for NeuN, Aβ42, Iba1 and GFAP Analysis

Mounted anti-NeuN, anti-Aβ42, anti-Iba1 and anti-GFAP stained sections were imaged on the Ti2 wide-field microscope (Northwestern University Center for Advanced Microscopy and Nikon Imaging Centre). Nikon NIS-Elements Software (Northwestern University Nikon Imaging Centre) was used to set the image intensity and size thresholds and was also used to manually trace the area of interest (hippocampus, cortex, or cerebellum). A binary channel was created for each area of interest for running the analysis.

### Antibodies

#### Antibodies used for immunohistochemistry

beta Amyloid Recombinant Rabbit Monoclonal Antibody (H31L21) (#700254, Thermo Fisher Scientific), Anti-GFAP antibody (#ab4674, Abcam), Anti-Iba1 antibody (#ab107159, Abcam), Anti-NeuN Antibody (#ABN91, EMD Millipore), and Alexa fluor-labeled secondary antibodies (Thermo Fisher Scientific).

#### Antibodies used for immunoblotting assay

Recombinant Anti-Angiotensin Converting Enzyme 1 antibody [EPR22291-247] (#ab254222, Abcam), Recombinant Anti-Angiotensin Converting Enzyme 1 antibody [EPR2757] (#ab75762, Abcam), Angiotensinogen Antibody (#79299, Cell Signaling Technology), Recombinant Anti-Angiotensin II Type 1 Receptor antibody [EPR3873] (#ab124734, Abcam), GAPDH (14C10) Rabbit mAb (#2118, Cell Signaling Technology), β-Actin Rabbit Monoclonal Antibody (#926-42210, LI-COR), Horse Anti-Mouse IgG Antibody (H+L), Peroxidase (#PI-2000-1, Vector Laboratories), and Goat Anti-Rabbit IgG Antibody (H+L), Peroxidase (#PI-1000-1, Vector Laboratories).

### Statistical Analysis

GraphPad Prism Version 9.4.1 (http://graphpad.com/scientific-software/prism/) was used for all statistical analysis. All data are presented as means ± SEM. Statistical significances were calculated by unpaired Student’s *t* test or ANOVA followed by Tukey’s post hoc test as detailed in the figure legends. *P* values less than 0.05 were considered significant: \**P* < 0.05, \*\**P* < 0.01, *** *P* < 0.001, and \*\*\*\**P* < 0.0001.

## Results

### Genetic knockdown of ACE1 in 5XFAD mice reduces ACE1 levels, while enalapril does not affect AngII levels in the brain

Previous study reported accelerated cerebral Aβ accumulation in 5XFAD mice that express five familial AD mutations associated with amyloid precursor protein (APP) and presenilin1 (PS1)^4^. In these mice, amyloid deposition initiated at around 2-months of age and rapidly accumulated overtime^4^. Furthermore, these mice displayed characteristic AD phenotypes, such as spatial memory deficits, gliosis, and neurodegeneration^4^. In this study, we sought to investigate the impact of ACE1 loss of function on amyloid burden by analyzing 6-month-old 5XFAD mice. ACE1 loss of function was mediated either by crossing 5XFAD mice to ACE1 conditional knockout mice or treating 5XFAD mice with the ACE1 inhibitor enalapril.

First, to examine 5XFAD mice with ACE1 genetic knockdown, we crossed 5XFAD mice with ACE1^FL/FL^iCre mice to generate 5XFAD; ACE1^FL/FL^iCre mice. To confirm ACE1 genetic knockout in 5XFAD; ACE1^FL/FL^iCre mice, we analyzed hippocampal and cortical homogenates of 5XFAD; ACE1^+/+^iCre mice and 5XFAD; ACE1^FL/FL^ control mice. Compared to controls, 5XFAD; ACE1^FL/FL^iCre mice showed around 63% and 50-58% reduction in ACE1 levels at the hippocampus and cortex respectively, indicating ablation of ACE1 in the excitatory forebrain neurons of 5XFAD; ACE1^FL/FL^iCre mice (Fig. 1A-D).

**Fig. 1.**
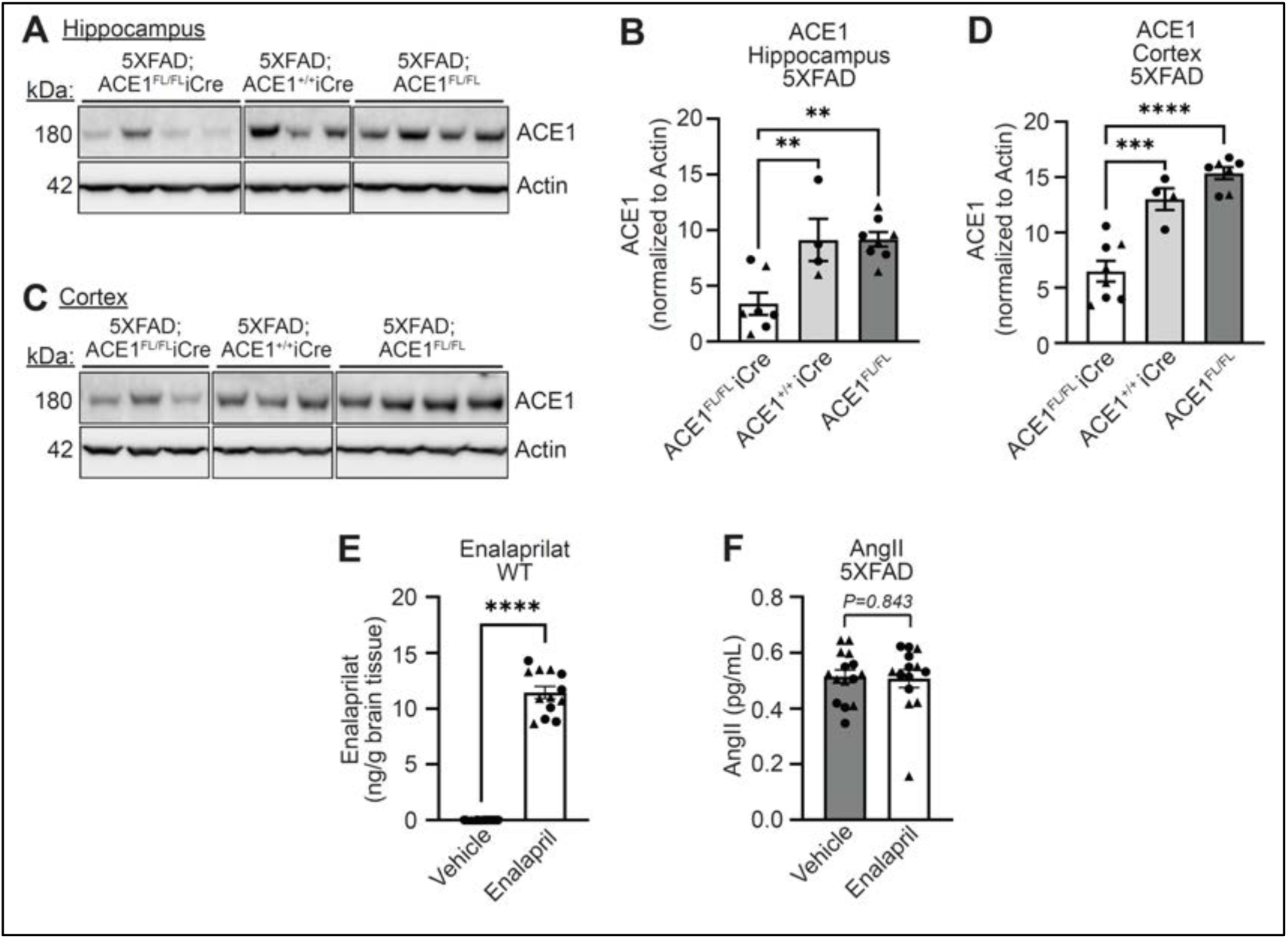
Genetic knockdown of ACE1 in 5XFAD mice reduces ACE1 levels, while enalapril does not affect AngII levels in the brain. **(A)** Immunoblot of hippocampal homogenates from 6-months old 5XFAD; ACE1^FL/FL^iCre, 5XFAD; ACE1^+/+^iCre, and 5XFAD; ACE1^FL/FL^ mice probed for ACE1 and Actin. Uncropped blots in Supplemental Figure 1A. **(B)** Quantification of ACE1 in (A) normalized to Actin (5XFAD; ACE1^FL/FL^iCre, n=7; 5XFAD; ACE1^+/+^iCre, n=4; 5XFAD; ACE1^FL/FL^, n=8). **(C)** Immunoblot of cortical homogenates from 6-months old 5XFAD; ACE1^FL/FL^iCre, 5XFAD; ACE1^+/+^iCre, and 5XFAD; ACE1^FL/FL^ mice probed for ACE1 and Actin. Uncropped blots in Supplemental Figure 1B. **(D)** Quantification of ACE1 in (C) normalized to Actin (5XFAD; ACE1^FL/FL^iCre, n=8; 5XFAD; ACE1^+/+^iCre, n=4; 5XFAD; ACE1^FL/FL^, n=7). **(E)** Measurement of Enalaprilat levels through liquid chromatography mass spectrometry (LC-MS) analysis by Pharmaron (Exton) Lab Services LLC. Quantification shows amount (ng) of drug per brain tissue (g) isolated from WT mice treated with vehicle or enalapril. **(F)** Quantification of AngII levels as measured by ELISA in vehicle or enalapril treated 5XFAD mice at 6-months of age (Vehicle treated 5XFAD, n=15; Enalapril treated 5XFAD, n=14). One-way ANOVA with Tukey’s multiple comparisons post hoc test in (B) and (D). Unpaired *t* test in (E) and (F). Circles represent females and triangles represent males. *P<0.05, **P<0.01, ***P<0.001, ****P<0.0001.

In parallel to the genetic knockdown, we investigated the effect of ACE1 pharmacological inhibition to cerebral amyloid accumulation. Specifically, wild-type (WT) and 5XFAD mice were administered enalapril, which is an antihypertensive drug that inhibits ACE1^45^. Enalapril is a prodrug that is hydrolyzed to enalaprilat for direct interaction with ACE1^26,34,45^. To examine enalapril delivery to the brain, we first measured the concentration of the active agent, enalaprilat, in the cerebellum of WT mice using liquid chromatography-tandem mass spectrometry (LC-MS/MS) analytical methods. We observed that enalapril treated mice showed significantly higher enalaprilat concentration compared to vehicle treated mice (Fig. 1E). In correlation, since ACE1 converts AngI to AngII, we next measured AngII levels in the mouse hemibrain homogenates by ELISA. We detected no significant differences in AngII levels in enalapril treated 5XFAD mice compared to vehicle treated 5XFAD mice in the brain (Fig. 1F). Together, our initial analysis confirmed genetic knockdown of ACE1 in 5XFAD; ACE1^FL/FL^iCre mice. Our results also show that while enalapril crossed the BBB in WT mice, AngII levels remained unchanged in enalapril treated 5XFAD mice compared to controls.

### ACE1 inhibition does not significantly alter the brain weight, body weight and brain region volumes of 5XFAD mice

In ACE1 KI mice with elevated ACE1 levels, there was significant reduction in hippocampal volume at 14-months of age, but not at 8-months of age^10^. This age-dependent hippocampal volume loss was rescued with captopril, an ACE1 inhibitor (ACEi), treatment^10^. To determine the neuronal and physiological effects of ACE1 inhibition in 5XFAD mice, we measured the brain and body weights of 5XFAD; ACE1^FL/FL^ mice compared to 5XFAD; ACE1^FL/FL^iCre, and of 5XFAD mice treated with vehicle or enalapril. Compared to 5XFAD; ACE1^FL/FL^ mice, 5XFAD; ACE1^FL/FL^iCre mice showed no significant changes in both brain and body weights (Fig. 2A, S2A). Similarly, our analysis of 5XFAD mice showed no significant differences in the brain and body weights upon vehicle or enalapril administration (Fig. 2B, S2B). We subsequently performed brain region volume assessment through measurements of brain sections from bregma coordinates ranging from approximately −0.94 to −3.40mm. Volume analysis revealed no significant differences in the hippocampal, cortical, and cerebellar volumes in 5XFAD mice treated with vehicle or enalapril, further reflecting our brain weight data (Fig. 2C-F). Additional analysis of enalapril treated WT mice mirrored that of 5XFAD mice (Fig. S2C-F). Therefore, these results suggest that ACE1 inhibition does not cause changes in hippocampal, cortical or cerebellar volumes in 5XFAD mice.

**Fig. 2.**
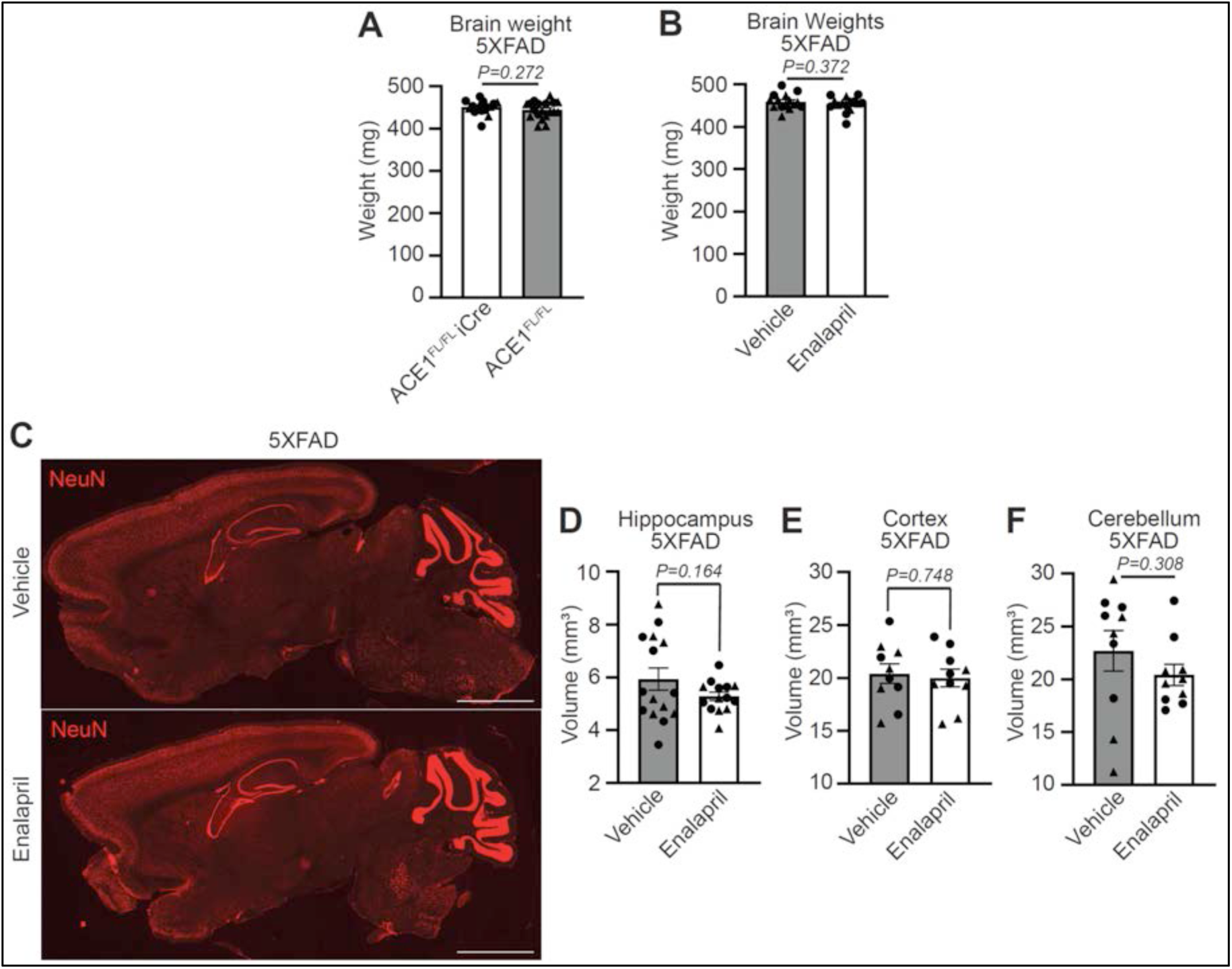
ACE1 inhibition does not significantly alter the brain weight and brain region volume of 5XFAD mice. **(A)** Quantification of the brain weights (mg) from 6-months old 5XFAD; ACE1^FL/FL^iCre and 5XFAD; ACE1^FL/FL^ mice. (5XFAD; ACE1^FL/FL^iCre, n=18; 5XFAD; ACE1^FL/FL^, n=19). **(B)** Quantification of the brain weights (mg) of vehicle or enalapril treated 5XFAD mice. (Vehicle treated 5XFAD, n=15; enalapril treated 5XFAD, n=15). **(C)** Representative images of sagittal brain sections from 6-months-old vehicle or enalapril treated 5XFAD mice. Sections were immunostained for NeuN (red) and imaged by confocal immunofluorescence microscopy. Scale bar, 2000μm. **(D)** Quantification of hippocampal volume in 5XFAD mice treated with vehicle or enalapril in (C). (Vehicle treated 5XFAD, n=15; Enalapril treated 5XFAD, n=14). **(E)** Quantification for cortical volume in 5XFAD mice treated with vehicle or enalapril in (C). (Vehicle treated 5XFAD, n=10; Enalapril treated 5XFAD, n=10). **(F)** Quantification for cerebellar volume in 5XFAD mice treated with vehicle or enalapril in (C). (Vehicle treated 5XFAD, n=10; Enalapril treated 5XFAD, n=10). Unpaired *t* test in (A to B) and (D to F). Circles represent females and triangles represent males. *P<0.05, **P<0.01, ***P<0.001, ****P<0.0001.

### Genetic and pharmacological ACE1 inhibition does not affect cerebral amyloid plaque load in the hippocampus and cortex of 5XFAD mice

Although multiple *in vitro* studies support ACE1’s role in Aβ degradation, clinical and *in vivo* studies report conflicting results^17–26,28–32,37–39,42^. To interrogate whether ACE1 catalyzes Aβ and affects amyloid accumulation *in vivo*, we examined amyloid plaque load as measured by Aβ42 levels through immunofluorescence microscopy in the hippocampus and the cortex. Since previous studies on 5XFAD mice reported that amyloid plaques first appear in the deep layers of the cortex, we first examined the cortical brain regions for potential changes in plaque accumulation at 6-months of age^4^. We found no significant differences in the plaque covered area, size and count between 5XFAD; ACE1^FL/FL^iCre, 5XFAD; ACE1^+/+^iCre, and 5XFAD; ACE1^FL/FL^ mice in the cortex (Fig. 3A-D, S5A-C). Corresponding analysis of 5XFAD mice with vehicle or enalapril treatment also showed no statistically significant changes (Fig. 3E-H, S5D-F). These results suggest that ACE1 loss of function does not affect amyloid plaque accumulation in the cortex of 5XFAD mice at 6-months of age.

**Fig. 3.**
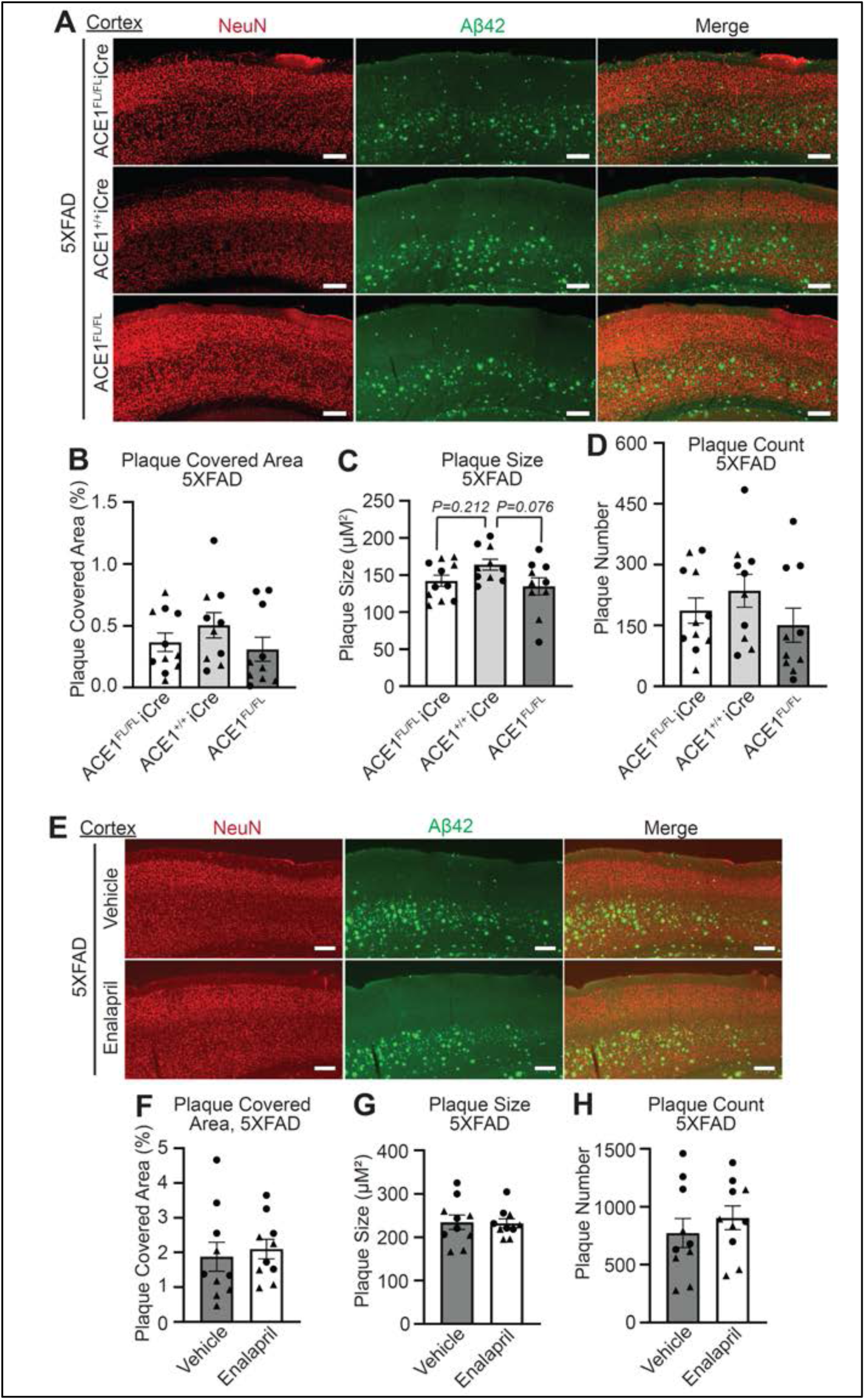
Both ACE1 knockout and inhibition does not affect cortical amyloid plaque load as examined by Aβ42 levels in 5XFAD mice. **(A)** Representative images showing layer 5 cortex of coronal sections from 6-months-old 5XFAD; ACE1^FL/FL^iCre, 5XFAD; ACE1^+/+^iCre, and 5XFAD; ACE1^FL/FL^ mice immunostained for NeuN (red) and Aβ42 (green) imaged by confocal immunofluorescence microscopy. Scale bar, 200μm. Quantification of plaque covered area **(B)**, plaque size **(C)**, and plaque count **(D)** in (A). (5XFAD; ACE1^FL/FL^iCre, n=11; 5XFAD; ACE1^+/+^iCre, n=10; 5XFAD; ACE1^FL/FL^, n=10). **(E)** Representative images showing layer 5 cortex of sagittal sections from 6-months-old vehicle and enalapril treated 5XFAD mice immunostained for NeuN (red) and Aβ42 (green) imaged by confocal immunofluorescence microscopy. Scale bar, 200μm. Quantification of plaque covered area **(F)**, plaque size **(G)**, and plaque count **(H)** in (E). (Vehicle treated 5XFAD, n=10; Enalapril treated 5XFAD, n=10). One-way ANOVA with Tukey’s multiple comparisons post hoc test was performed in (B to D). Unpaired *t* test in (F to H). Circles represent females and triangles represent males. *P<0.05, **P<0.01, ***P<0.001, ****P<0.0001.

In addition to the cortical brain regions, we further examined the hippocampal brain regions, which is another vulnerable brain region in AD. Recent study on mice with ACE1 conditional knockout in the excitatory forebrain neurons displayed hippocampal dependent behavioral deficits and showed RAS dysregulation and cerebrovascular defects selectively in the hippocampal brain regions. Moreover, an aforementioned study reported that ACE1 KI mice displayed hippocampal neurodegeneration which was rescued by ACEi administration^10^. These studies point to the impact of ACE1 selectively on the hippocampus in the AD brain and prompted us to investigate whether ACE1 affects hippocampal amyloid plaque accumulation. Our analysis of plaque covered area, size and count revealed no significant differences between 5XFAD; ACE1^FL/FL^iCre, 5XFAD; ACE1^+/+^iCre and 5XFAD; ACE1^FL/FL^ mice in the hippocampus (Fig. 4A-D, S6A-C). Similarly, assessment of plaque overed area, size, and count in enalapril treated 5XFAD mice also showed no significant differences in hippocampus compared to that in vehicle treated 5XFAD mice (Fig. 4E-H, S6D-F). These analyses suggest that ACE1 does not affect hippocampal amyloid plaque accumulation.

**Fig. 4.**
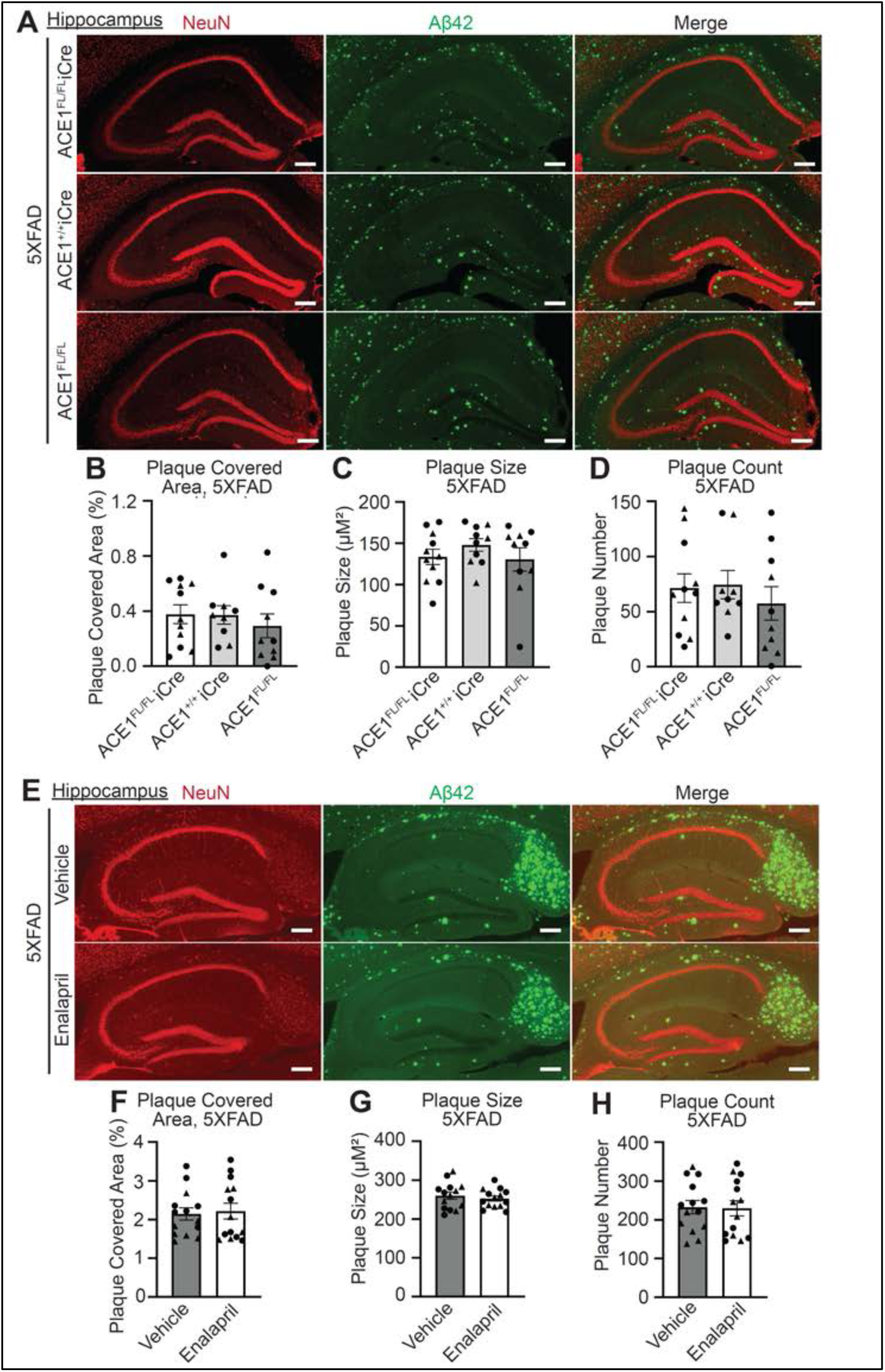
Both ACE1 genetic knockout and inhibition does not affect hippocampal amyloid plaque load as examined by Aβ42 levels in 5XFAD mice. **(A)** Representative images showing the hippocampus of coronal sections from 6-months-old 5XFAD; ACE1^FL/FL^iCre, 5XFAD; ACE1^+/+^iCre, and 5XFAD; ACE1^FL/FL^ mice immunostained for NeuN (red) and Aβ42 (green) imaged by confocal immunofluorescence microscopy. Scale bar, 200μm. Quantification of plaque covered area **(B)**, plaque size **(C)**, and plaque count **(D)** in (A). (5XFAD; ACE1^FL/FL^iCre, n=11; 5XFAD; ACE1^+/+^iCre, n=9-10; 5XFAD; ACE1^FL/FL^, n=10). **(E)** Representative images showing the hippocampus of sagittal sections from 6-months-old vehicle and enalapril treated 5XFAD mice immunostained for NeuN (red) and Aβ42 (green) imaged by confocal immunofluorescence microscopy. Scale bar, 200μm. Quantification of plaque covered area **(F)**, plaque size **(G)**, and plaque count **(H)** in (E). (Vehicle treated 5XFAD, n=14; Enalapril treated 5XFAD, n=14). One-way ANOVA with Tukey’s multiple comparisons post hoc test was performed in (B to D). Unpaired *t* test in (F to H). Circles represent females and triangles represent males. *P<0.05, **P<0.01, ***P<0.001, ****P<0.0001.

We also examine amyloid plaque accumulation in the cerebellum where amyloid plaques typically appear at later advanced stages in AD. As expected, cerebellar amyloid plaque levels were not significantly different between vehicle and enalapril treated 5XFAD mice at 6-months of age (Fig. S4A-D). Altogether, our results suggest that ACE1 does not affect amyloid plaque accumulation in 5XFAD mice *in vivo*.

### ACE1 inhibition does not affect neuroinflammation in the hippocampus and cortex of 5XFAD mice

Multiple studies report neuroinflammation in AD transgenic mouse models, including 5XFAD mice wherein astrocytes were closely associated with amyloid deposits in the brain^4^. ACE1 KI mice also showed neuroinflammation, which was accelerated in 5XFAD background but attenuated with ACEi administration^10^. We next explored whether ACE1 inhibition in 5XFAD mice impacts neuroinflammation in the brain. To investigate this, we examined glial fibrillary acidic protein (GFAP) and ionized calcium binding adaptor molecule 1 (Iba1) immunoreactivity as a measure for astrocytic and microglial activation.

We first examined the hippocampal brain regions where astrogliosis and microgliosis have been observed in both 5XFAD and ACE1 KI mice based on previous studies ^4,10^. Immunofluorescence microscopy analysis showed no statistically significant differences in the GFAP and Iba1 covered area in the hippocampus between 5XFAD; ACE1^FL/FL^iCre, 5XFAD; ACE1^+/+^iCre and 5XFAD; ACE1^FL/FL^ mice (Fig. 5A-C). Likewise, corresponding analysis of 5XFAD mice treated with enalapril or vehicle also showed that ACE1 inhibition does not affect neuroinflammation in 5XFAD mice (Fig. 5D-F). Thus, ACE1 loss of function in the hippocampus does not affect neuroinflammation in 5XFAD mice.

**Fig. 5.**
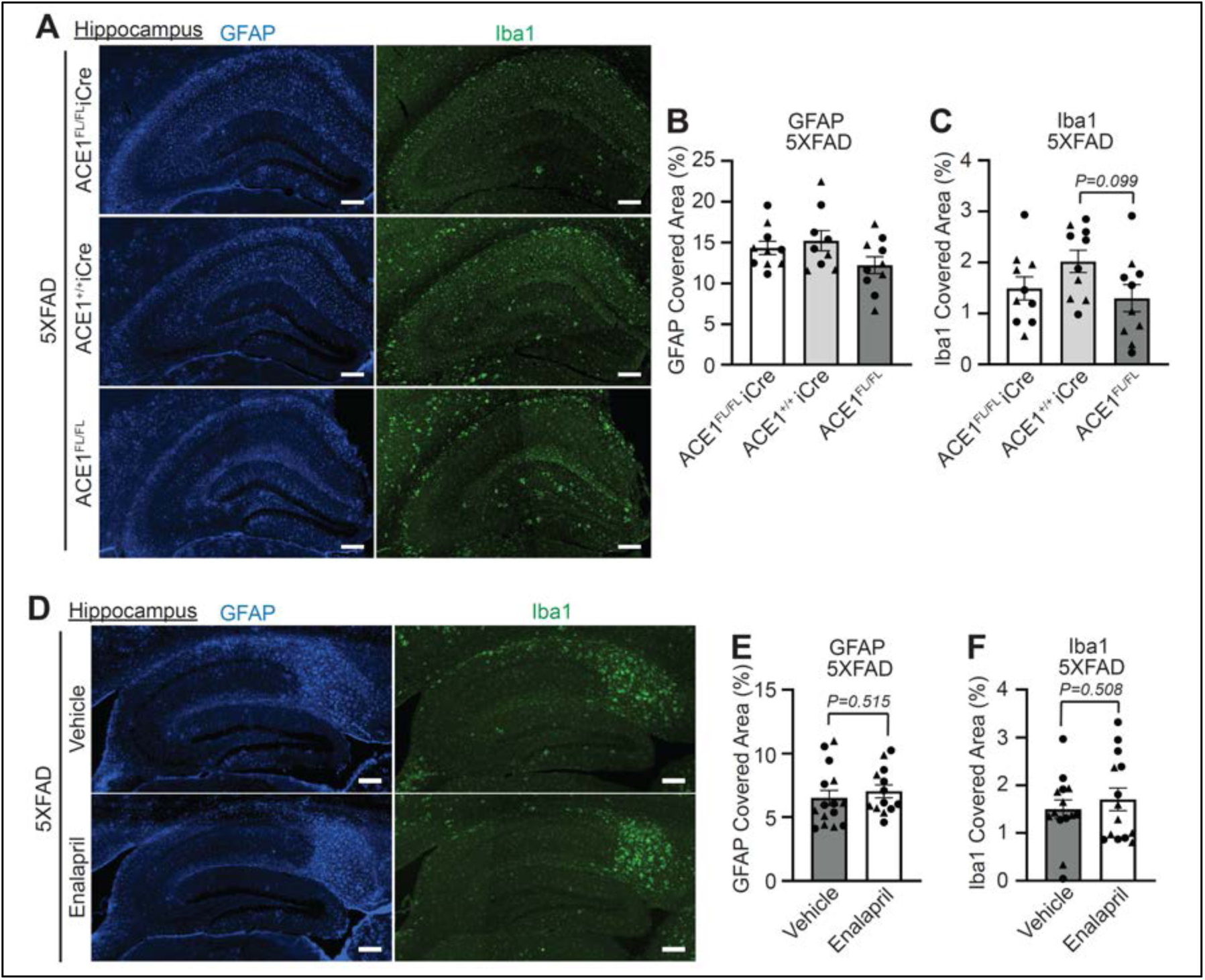
Hippocampal neuroinflammation is unaffected in 5XFAD mice with ACE1 genetic knockout or enalapril treatment. **(A)** Representative images showing the hippocampus of coronal sections from 6-months-old 5XFAD; ACE1^FL/FL^iCre, 5XFAD; ACE1^+/+^iCre, and 5XFAD; ACE1^FL/FL^ mice immunostained for GFAP (blue) and Iba1 (green) imaged by confocal immunofluorescence microscopy. Scale bar, 200μm. Quantification of GFAP covered area **(B)** and Iba1 covered area **(C)** in (A). (5XFAD; ACE1^FL/FL^iCre, n=10; 5XFAD; ACE1^+/+^iCre, n=9-10; 5XFAD; ACE1^FL/FL^, n=10). **(D)** Representative images showing the hippocampus of sagittal sections from 6-months-old vehicle and enalapril treated 5XFAD mice immunostained for GFAP (blue) and Iba1 (green) imaged by confocal immunofluorescence microscopy. Scale bar, 200μm. Quantification of GFAP covered area **(E)** and Iba1 covered area **(F)** in (D). (Vehicle treated 5XFAD, n=14-15; Enalapril treated 5XFAD, n=13-14). One-way ANOVA with Tukey’s multiple comparisons post hoc test was performed in (B) and (C). Unpaired *t* test in (E) and (F). Circles represent females and triangles represent males. *P<0.05, **P<0.01, ***P<0.001, ****P<0.0001.

Robust gliosis has been also observed in the cortical brain regions in both 5XFAD with and without ACE1 KI mutations^4,10^. Our analysis did not detect significant changes in astrogliosis and microgliosis between 5XFAD; ACE1^FL/FL^iCre, 5XFAD; ACE1^+/+^iCre and 5XFAD; ACE1^FL/FL^ mice as measured by GFAP and Iba1 coverage (Fig. 6A-C). In line with this, enalapril or vehicle treated 5XFAD mice also did not show any differences in neuroinflammation (Fig. 6D-F). Additional analysis of cerebellar brain regions also did not show any signs of neuroinflammation (Fig. S4E-G). Together, in 5XFAD mice, neuroinflammation is unaffected with ACE1 loss of function in the cortical brain regions.

**Fig. 6.**
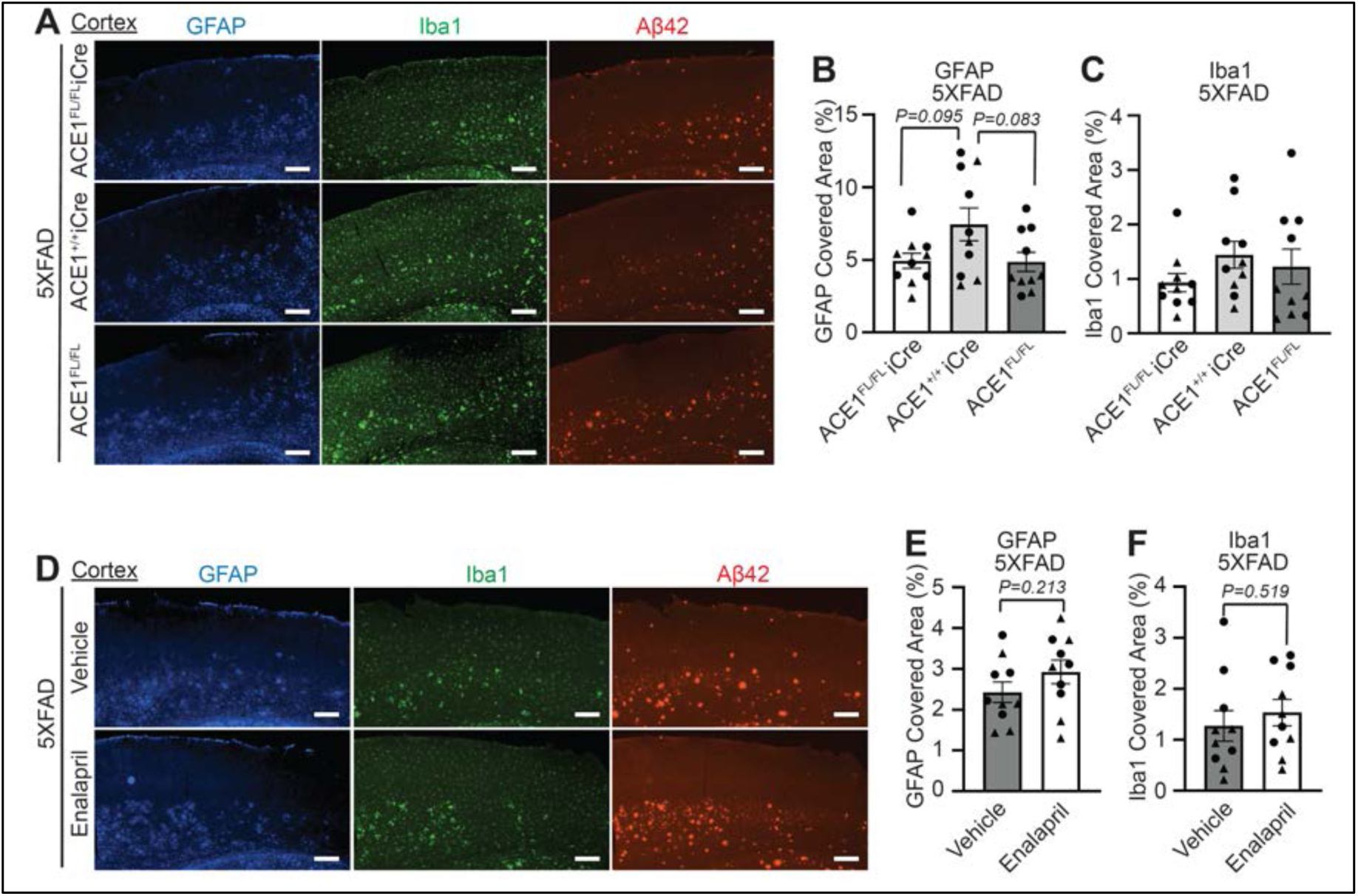
Cortical neuroinflammation is unaffected in 5XFAD mice with ACE1 genetic knockout or enalapril treatment. **(A)** Representative images showing layer 5 cortex of coronal sections from 6-months-old 5XFAD; ACE1^FL/FL^iCre, 5XFAD; ACE1^+/+^iCre, and 5XFAD; ACE1^FL/FL^ mice immunostained for GFAP (blue), Iba1 (green) and Aβ42 (red) imaged by confocal immunofluorescence microscopy. Scale bar, 200μm. Quantification of GFAP covered area **(B)** and Iba1 covered area **(C)** in (A). (5XFAD; ACE1^FL/FL^iCre, n=10-11; 5XFAD; ACE1^+/+^iCre, n=10; 5XFAD; ACE1^FL/FL^, n=10). **(D)** Representative images showing layer 5 cortex of sagittal sections from 6-months-old vehicle and enalapril treated 5XFAD mice immunostained for GFAP (blue), Iba1 (green) and Aβ42 (red) imaged by confocal immunofluorescence microscopy. Scale bar, 200μm. Quantification of GFAP covered area **(E)** and Iba1 covered area **(F)** in (D). (Vehicle treated 5XFAD, n=10; Enalapril treated 5XFAD, n=10). One-way ANOVA with Tukey’s multiple comparisons post hoc test was performed in (B) and (C). Unpaired *t* test in (E) and (F). Circles represent females and triangles represent males. *P<0.05, **P<0.01, ***P<0.001, ****P<0.0001.

## Discussion

In this study, we interrogated the discrepancies regarding the role of ACE1 in Aβ accumulation. We examined this *in vivo* in a novel 5XFAD ACE1 conditional knockout mouse model. Furthermore, we performed a pharmacological study in which 5XFAD mice were treated with a clinically relevant ACE1 inhibitor. Our results show that, in the 5XFAD mouse model, ACE1 inhibition does not alter Aβ levels, suggesting that ACE1 does not play a significant role in Aβ degradation *in vivo*.

The major finding in this study is that ACE1 inhibition in the transgenic AD brain does not cause an alteration in amyloid burden. This conflicts with most of the existing *in vitro* studies reporting ACE1’s role in Aβ degradation and thus implies the important role of ACE1 to mitigate AD pathogenesis^18–24^. In particular, a research conducted on neuroblastoma cells reported that enalaprilat inhibits zinc-bound Aβ(1-16) dimer, which may be a neurotoxic form in AD^46^. Our findings also conflict with multiple *in vivo* and clinical studies suggesting that ACE1 exacerbates AD pathogenesis; therefore, emphasizing the benefits of ACE1 inhibition^28–33,35,36,42^. In contrast, our results are in line with several *in vivo* and clinical studies suggesting that ACE1 does not affect Aβ levels in the brain^23,37–39,41–44^. Specifically, several studies on AD transgenic mice revealed no significant differences in cerebral Aβ levels with ACE1 inhibitors^37–40^. In addition, multiple lines of evidence from clinical studies report no significant differences in AD incidence with ACE1 inhibitor as a class^35,42,43^. Furthermore, investigation of AD transgenic mice with AT1R null mutation showed reduced amyloid pathology, but ACE1 levels remained unaltered^41^. Therefore, our findings substantiate some clinical and *in vivo* studies but not *in vitro* reports.

Our physiological and functional studies demonstrate the ACE1 inhibition in 5XFAD mice does not cause neuroinflammation and brain atrophy, which was unexpected (Fig. 2-4, S4B). Based on the canonical RAS pathway, ACE1 inhibition should reduce AngII level and subsequently inhibit AT1R activation. Given that AT1R activation causes increased neuroinflammation, oxidative stress and cell death, we expected to see rescue in neuroinflammation and brain atrophy upon reduction in AT1R activation through ACE1 inhibition^17,47^. Our expectation is also based on previous study where AD-induced rat treated with BBB-penetrating ACE1 inhibitor, perindopril, showed attenuation in neuroinflammation^48^. The observed unexpected result is likely due to the unaltered AngII levels in enalapril treated 5XFAD mice, potentially by compensatory mechanisms (Fig. 1F). For example, in the noncanonical RAS pathway, cathepsin G or tonin can directly produce AngII from angiotensinogen (AGT)^12^. Thus, this unaltered AT1R activation through compensated AngII levels may have led to no rescue of neuroinflammation and brain atrophy as observed in our data.

## Conclusion

In conclusion, ACE1 does not affect plaque accumulation in 5XFAD transgenic AD mouse model. Therefore, the mechanism of ACE1 mutation in AD pathogenesis may be through alternative pathways that remains to be explored in future studies. Furthermore, our data implies that antihypertensive ACE1 inhibitors, such as enalapril, does not directly ameliorate nor exacerbate cerebral plaque accumulation and neuroinflammation in mice. These findings may be of additional relevance to hypertensive patients under blood pressure medication and implies the need for further investigation to understand alternative causes for the cognitive changes observed in previous studies.

## Abbreviations

Aβ42: Amyloid Beta 42 peptide
ACE1: Angiotensin Converting Enzyme 1
ACEi: ACE1 inhibitor
AD: Alzheimer’s Disease
AGT: Angiotensinogen
AngI: Angiotensin I
AngII: Angiotensin II
APP: Amyloid Precursor Protein
AT1R: AngII Receptor Type 1
AT2R: AngII Receptor Type 2
BBB: Blood Brain Barrier
CER: Cerebellum
CNS: Central Nervous System
CTX: Cortex
FAD: Familial Alzheimer’s Disease
HPC: Hippocampus
LC-MS/MS: Liquid chromatography - tandem mass spectrometry
PS1: Presenilin1
RAS: Renin Angiotensin System

## Declarations

### Ethics approval and consent to participate

All animal work was performed in accordance with Northwestern University Institutional Animal Care and Use Committee approval.

## Consent for publication

Not applicable.

## Availability of data and materials

The datasets used and/or analyzed during the current study are available from the corresponding author on reasonable request.

## Competing interests

The authors declare that they have no competing interests.

## Funding

This study was supported by the Cure Alzheimer’s Fund and the National Institute on Aging (AG080092-01).

## Authors’ contributions

Conceptualization, LKC, RV. Methodology, validation, data analysis, and investigation, SJ, AA, LKC, RV. Interpretation, SJ, LKC, RV. Writing original draft, SJ. Editing, SJ, LKC, RV. Supervision, LKC, RV. Funding acquisition, RV. Visualization, SJ, LKC, RV.

## Acknowledgements

We thank all members of the Vassar laboratory for helpful suggestions. We thank the teams at the Northwestern University Center for Advanced Microscopy for expert advice on imaging analysis and the Northwestern University Center for Comparative Medicine for animal care support. We acknowledge M. Senft, D. Hidalgo, R. MacNeill, and R. Strab of Pharmaron (Exton) Lab Services LLC for the LC-MS/MS analysis on enalaprilat concentration measurements. Lastly, we are grateful to L. Pieramici for technical assistance.

## Figure and Figure Legends

**Supplemental Fig. 1.**
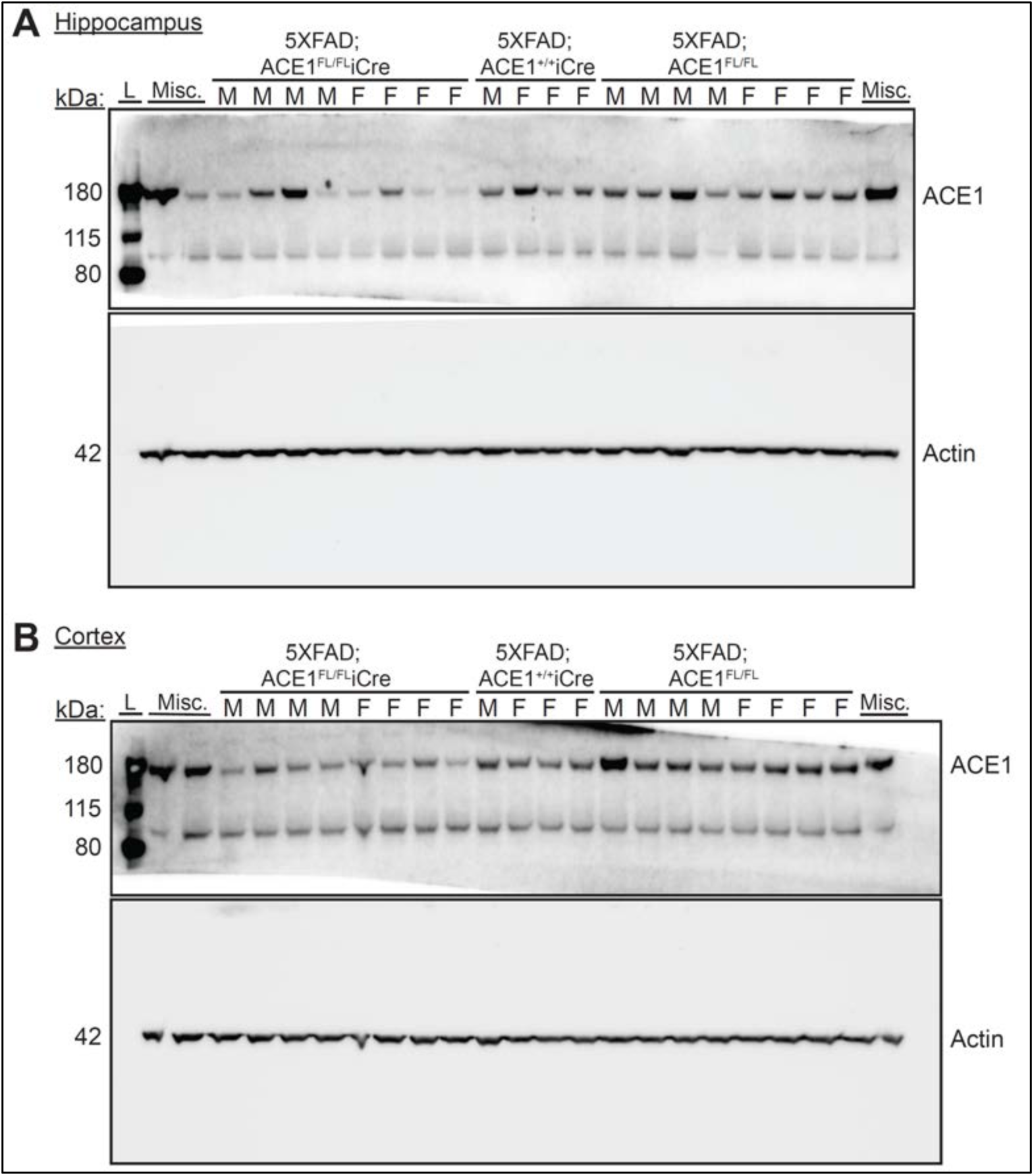
Original blots of ACE1 shown in Main Figure 1. **(A)** Original (uncropped) immunoblots (of Fig. 1A) showing hippocampal homogenates from 6-months old 5XFAD; ACE1^FL/FL^iCre, 5XFAD; ACE1^+/+^iCre, and 5XFAD; ACE1^FL/FL^ mice probed for ACE1 and Actin. (5XFAD; ACE1^FL/FL^iCre, n=7; 5XFAD; ACE1^+/+^iCre, n=4; 5XFAD; ACE1^FL/FL^, n=8). **(B)** Original immunoblots (of Fig. 1C) showing cortical homogenates from 6-months old 5XFAD; ACE1^FL/FL^iCre, 5XFAD; ACE1^+/+^iCre, and 5XFAD; ACE1^FL/FL^ mice probed for ACE1 and Actin. (5XFAD; ACE1^FL/FL^iCre, n=8; 5XFAD; ACE1^+/+^iCre, n=4; 5XFAD; ACE1^FL/FL^, n=7). Abbreviations (L = protein ladder; Misc. = miscellaneous sample; F = females; M = males).

**Supplemental Fig. 2.**
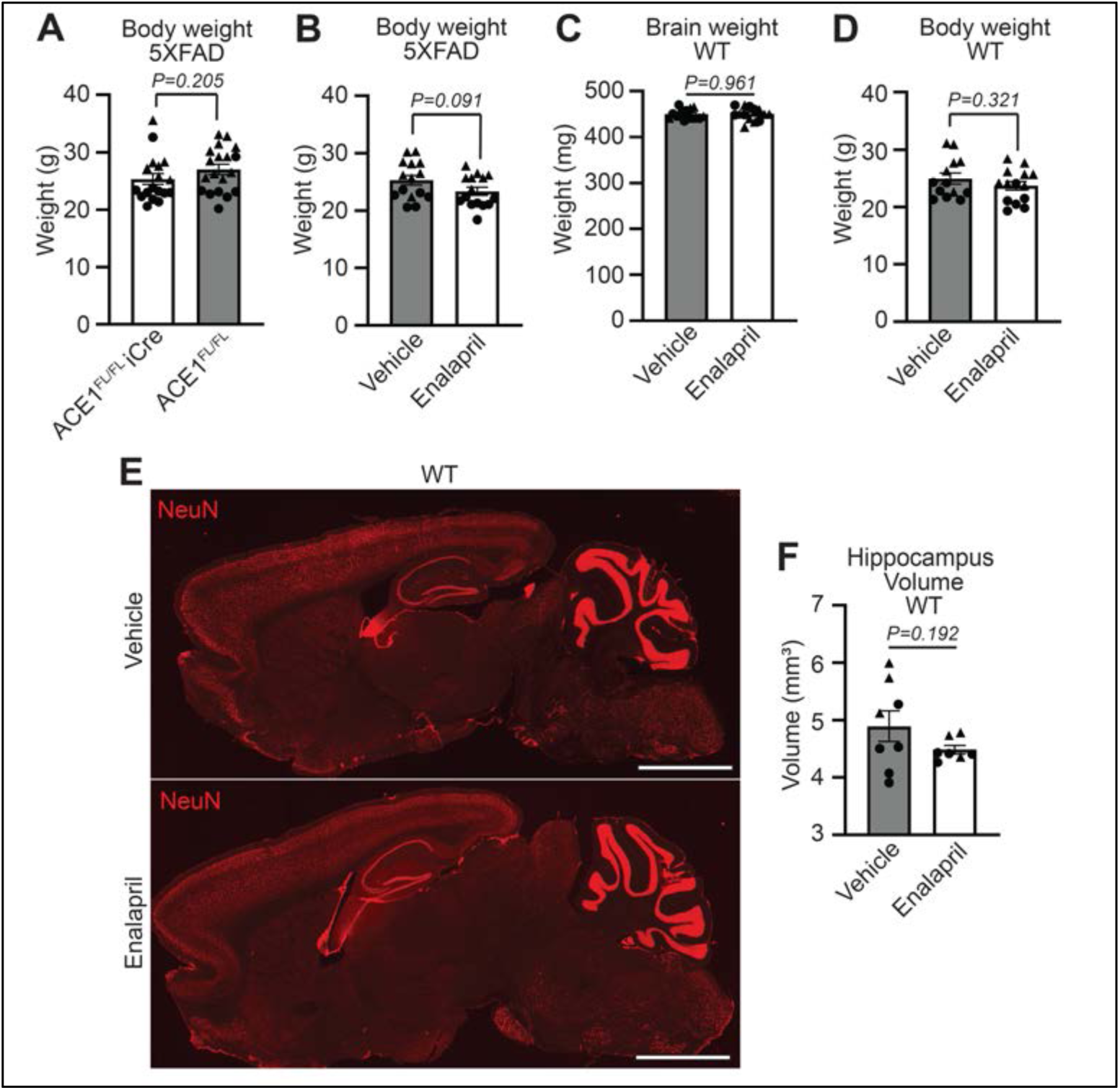
5XFAD mice show no changes in body weight and WT mice show no changes in brain weight, body weight, and hippocampal volume. **(A)** Quantification of the body weights (g) from 6-months old 5XFAD; ACE1^FL/FL^iCre and 5XFAD; ACE1^FL/FL^ mice (5XFAD; ACE1^FL/FL^iCre, n=18; 5XFAD; ACE1^FL/FL^, n=19). **(B)** Quantification of the body weights (g) of vehicle or enalapril treated 5XFAD mice. (Vehicle treated 5XFAD, n=15; enalapril treated 5XFAD, n=15). **(C)** Quantification of the brain weights (mg) of vehicle or enalapril treated wild-type (WT) mice. (Vehicle treated WT, n=15; enalapril treated WT, n=14). **(D)** Quantification of the body weights (g) of vehicle or enalapril treated WT mice. (Vehicle treated WT, n=13; enalapril treated WT, n=14). **(E)** Representative sagittal brain sections from 6-months-old vehicle or enalapril treated WT mice. Sections were immunostained for NeuN (red) and imaged by confocal immunofluorescence microscopy. Scale bar, 2000μm. **(F)** Quantification of hippocampal volume in WT mice treated with vehicle or enalapril in (E). (Vehicle treated WT, n=8; Enalapril treated WT, n=7). Unpaired *t* test in (A to D) and (F). Circles represent females and triangles represent males. *P<0.05, **P<0.01, ***P<0.001, ****P<0.0001.

**Supplemental Figure 3.**
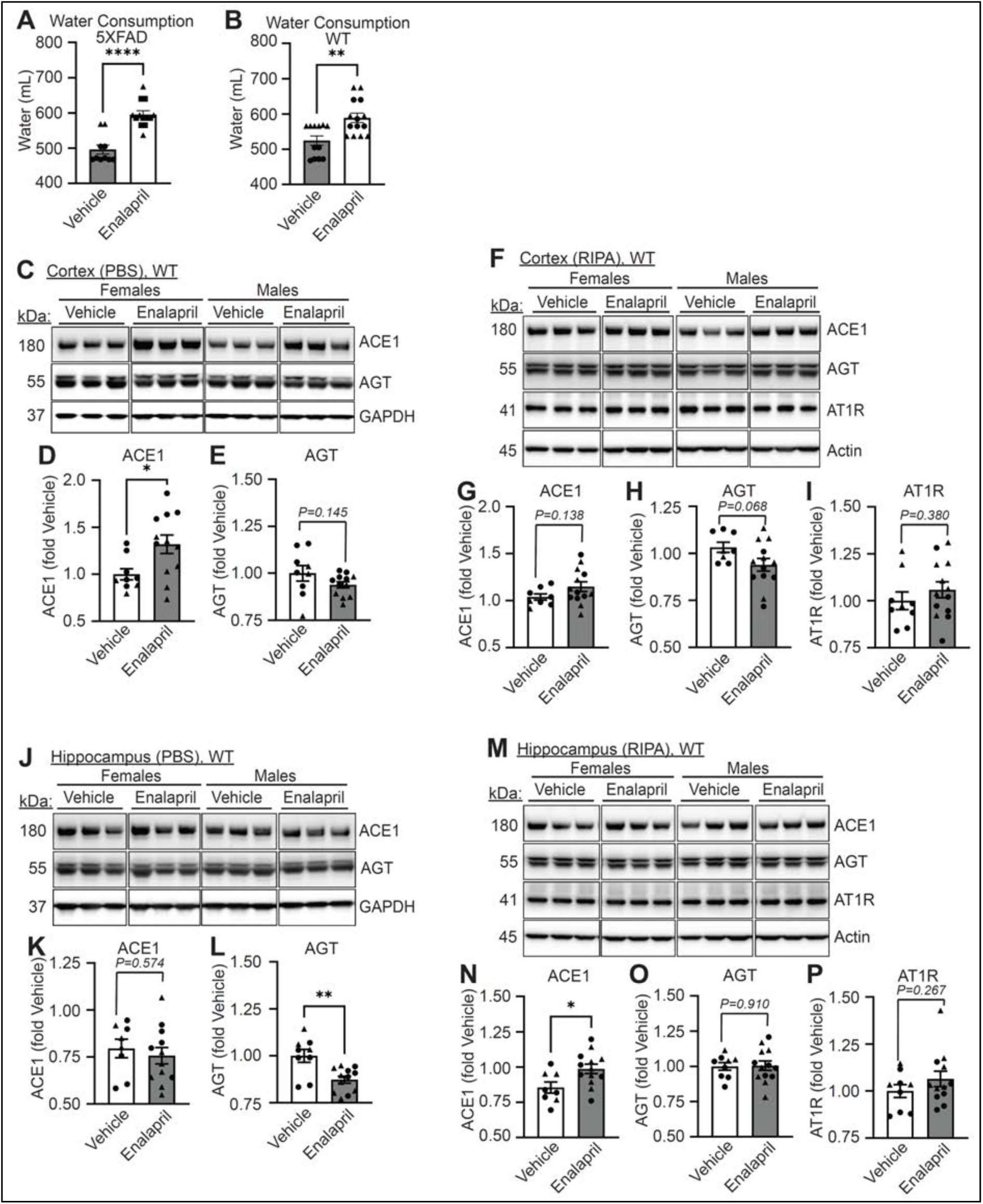
Increased water consumption and RAS component alterations with enalapril treatment. **(A)** Quantification of the amount of water (ml) consumed by vehicle or enalapril treated 5XFAD mice over 15-weeks. (Vehicle treated 5XFAD, n=10; enalapril treated 5XFAD, n=13). **(B)** Quantification of the amount of water (ml) consumed by vehicle or enalapril treated WT mice over 15-weeks (vehicle treated WT, n=12; enalapril treated WT, n=14). **(C)** Immunoblots of PBS-soluble cortical homogenates from 6-months-old WT mice treated with vehicle or enalapril that were probed for ACE1, Angiotensinogen (AGT), and GAPDH. Quantification of ACE1 **(D)** and AGT **(E)** in (C) normalized to GAPDH (Vehicle treated WT, n=9; Enalapril treated WT, n=12). **(F)** Immunoblots of RIPA-soluble cortical homogenates from 6-months-old WT mice treated with vehicle or enalapril that were probed for ACE1, AGT, AT1R, and Actin. Quantification of ACE1 **(G)**, AGT **(H)**, and AT1R **(I)** in (F) normalized to Actin (Vehicle treated WT, n=8-9; Enalapril treated WT, n=12-13). **(J)** Immunoblots of PBS-soluble hippocampal homogenates from 6-months-old WT mice treated with vehicle or enalapril that were probed for ACE1, AGT, and GAPDH. Quantification of ACE1 **(K)** and AGT **(L)** in (J) normalized to GAPDH (Vehicle treated WT, n=8-9; Enalapril treated WT, n=12). **(M)** Immunoblots of RIPA-soluble hippocampal homogenates from 6-months-old WT mice treated with vehicle or enalapril that were probed for ACE1, AGT, AT1R, and Actin. Quantification of ACE1 **(N)**, AGT **(O)**, and AT1R **(P)** in (M) normalized to Actin (Vehicle treated WT, n=8-9; Enalapril treated WT, n=12-13). Unpaired *t* test in (A and B), (D and E), (G to I), (K and L), and (N to P). Circles represent females and triangles represent males. *P<0.05, **P<0.01, ***P<0.001, ****P<0.0001.

**Supplemental Fig. 4.**
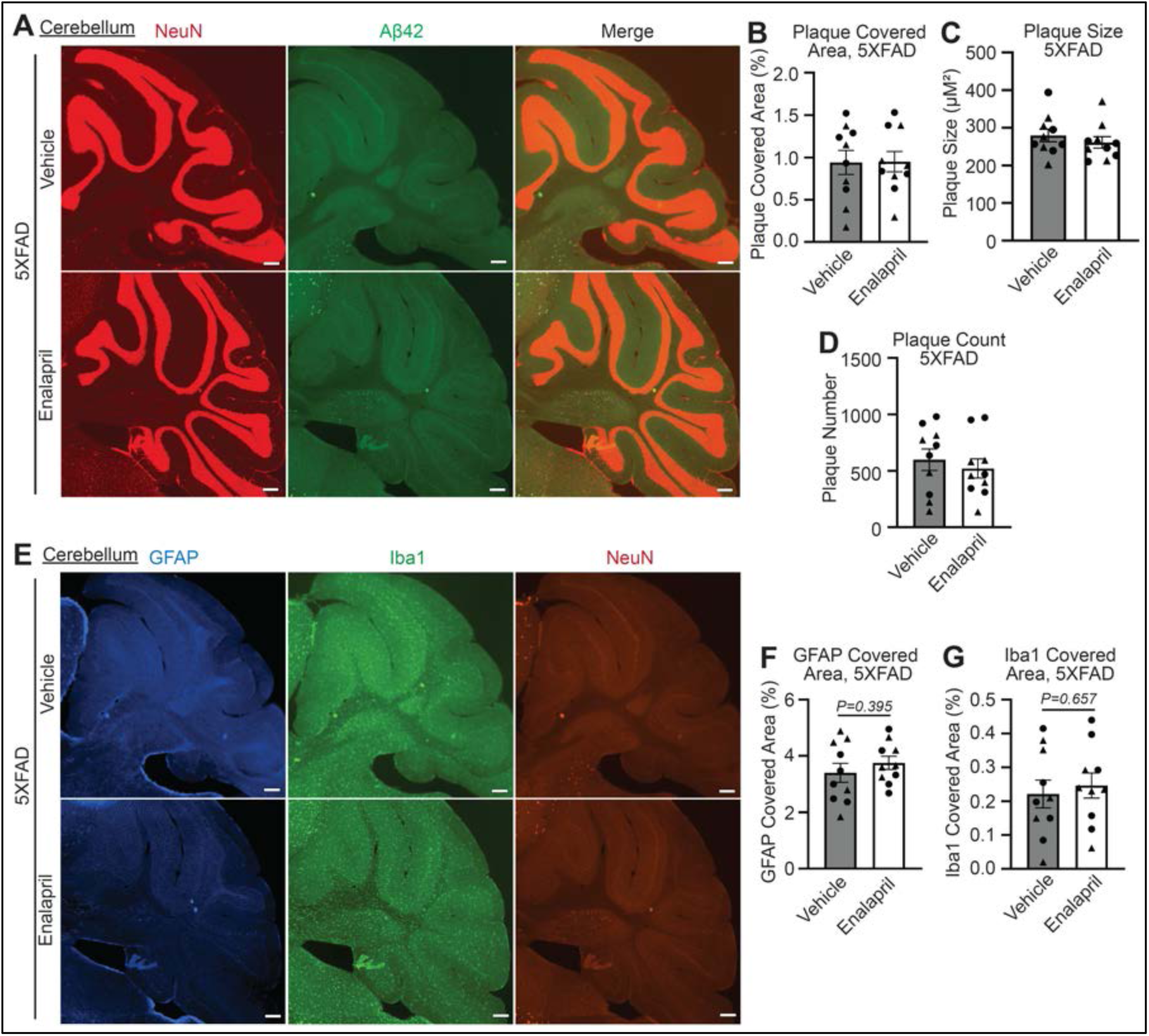
Amyloid plaque burden and neuroinflammation are unaffected in enalapril-treated 5XFAD mice in the cerebellum. **(A)** Representative images showing the cerebellum of sagittal sections from 6-months-old vehicle and enalapril treated 5XFAD mice immunostained for NeuN (red) and Aβ42 (green) imaged by confocal immunofluorescence microscopy. Scale bar, 200μm. Quantification of plaque covered area **(B)**, plaque size **(C)**, and plaque count **(D)** in (A). (Vehicle treated 5XFAD, n=10; Enalapril treated 5XFAD, n=10). **(E)** Representative images showing the cerebellum of sagittal sections from 6-months-old vehicle and enalapril treated 5XFAD mice immunostained for GFAP (blue) and Iba1 (green) imaged by confocal immunofluorescence microscopy. Scale bar, 200μm. Quantification of GFAP covered area **(F)** and Iba1 covered area **(G)** in (E). (Vehicle treated 5XFAD, n=9-10; Enalapril treated 5XFAD, n=9-10). Unpaired *t* test in (B to D), (F), and (G). Circles represent females and triangles represent males. *P<0.05, **P<0.01, ***P<0.001, ****P<0.0001.

**Supplemental Fig. 5.**
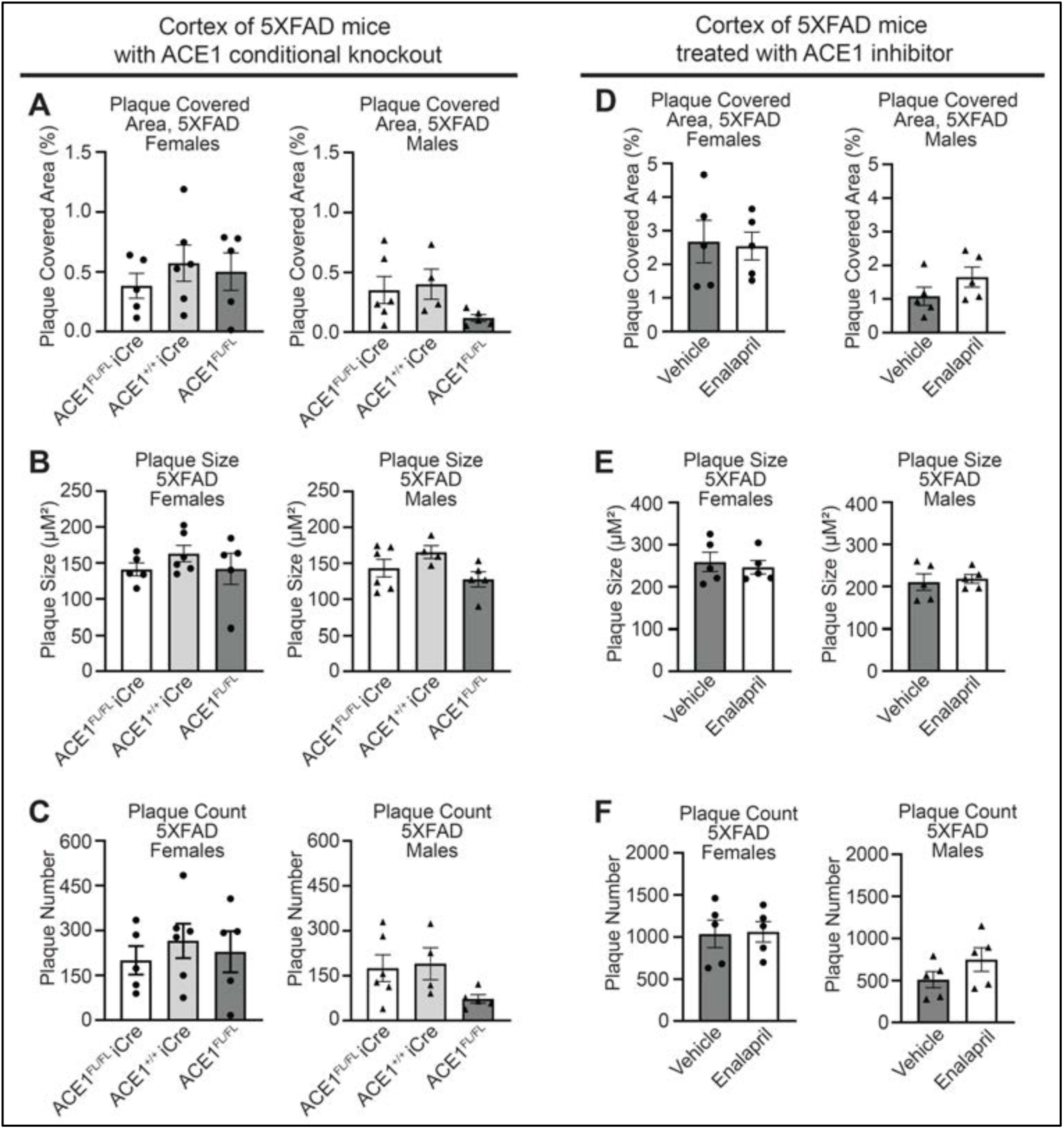
Cortical plaque covered area, size and count are unaltered in both 5XFAD males and females with ACE1 genetic knockdown or enalapril. **(A to C)** Imaging analysis of coronal sections from the cortex of 6-months-old 5XFAD; ACE1^FL/FL^iCre, 5XFAD; ACE1^+/+^iCre, and 5XFAD; ACE1^FL/FL^ mice. Quantification of plaque covered area **(A)**, plaque size **(B)**, and plaque count **(C)** in (Fig. 5A) through independent analysis of females (left) and males (right). <u>Females</u> (5XFAD; ACE1^FL/FL^iCre, n=5; 5XFAD; ACE1^+/+^iCre, n=6; 5XFAD; ACE1^FL/FL^, n=5). <u>Males</u> (5XFAD; ACE1^FL/FL^iCre, n=6; 5XFAD; ACE1^+/+^iCre, n=4; 5XFAD; ACE1^FL/FL^, n=5). **(D to F)** Imaging analysis of sagittal sections from the cortex of 6-months-old Vehicle or Enalapril treated 5XFAD mice. Quantification of plaque covered area **(D)**, plaque size **(E)**, and plaque count **(F)** in (Fig. 5E) through independent analysis of females (left) and males (right). <u>Females</u> (Vehicle treated 5XFAD, n=5; Enalapril treated 5XFAD, n=5). <u>Males</u> (Vehicle treated 5XFAD, n=5; Enalapril treated 5XFAD, n=5). One-way ANOVA with Tukey’s multiple comparisons post hoc test was performed in (A to C). Unpaired *t* test in (D to F). Circles represent females and triangles represent males. *P<0.05, **P<0.01, ***P<0.001, ****P<0.0001.

**Supplemental Fig. 6.**
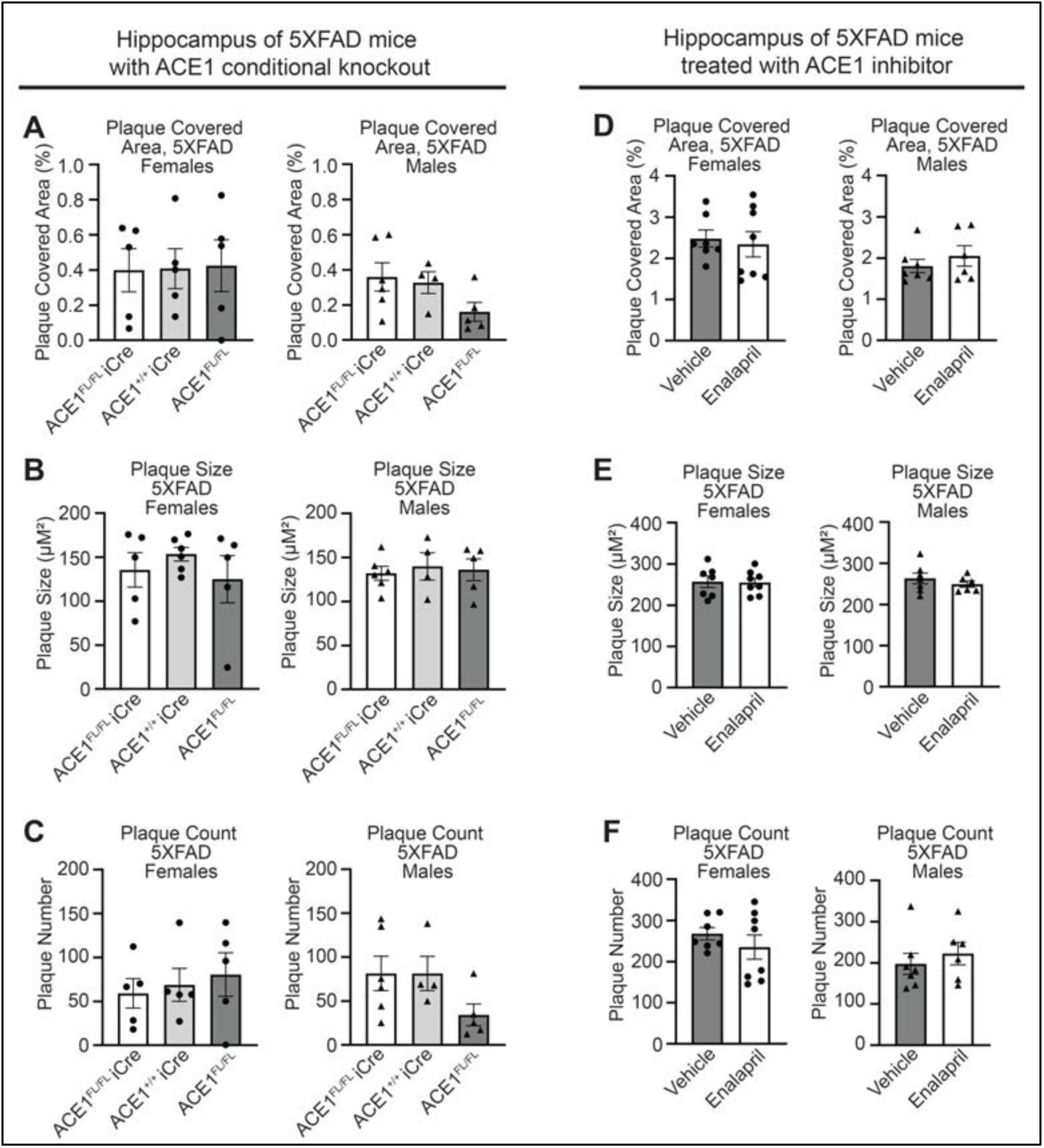
Hippocampal plaque covered area, size and count are unchanged in both 5XFAD males and females with ACE1 genetic knockdown or enalapril. **(A to C)** Imaging analysis of coronal sections from the hippocampus of 6-months-old 5XFAD; ACE1^FL/FL^iCre, 5XFAD; ACE1^+/+^iCre, and 5XFAD; ACE1^FL/FL^ mice. Quantification of plaque covered area **(A)**, plaque size **(B)**, and plaque count **(C)** in (Fig. 6A) through independent analysis of females (left) and males (right). <u>Females</u> (5XFAD; ACE1^FL/FL^iCre, n=5; 5XFAD; ACE1^+/+^iCre, n=5-6; 5XFAD; ACE1^FL/FL^, n=5). <u>Males</u> (5XFAD; ACE1^FL/FL^iCre, n=6; 5XFAD; ACE1^+/+^iCre, n=4; 5XFAD; ACE1^FL/FL^, n=5). **(D to F)** Imaging analysis of sagittal sections from the hippocampus of 6-months-old Vehicle or Enalapril treated 5XFAD mice. Quantification of plaque covered area **(D)**, plaque size **(E)**, and plaque count **(F)** in (Fig. 6E) through independent analysis of females (left) and males (right). <u>Females</u> (Vehicle treated 5XFAD, n=7; Enalapril treated 5XFAD, n=8). <u>Males</u> (Vehicle treated 5XFAD, n=7; Enalapril treated 5XFAD, n=6). One-way ANOVA with Tukey’s multiple comparisons post hoc test was performed in (A to C). Unpaired *t* test in (D to F). Circles represent females and triangles represent males. *P<0.05, **P<0.01, ***P<0.001, ****P<0.0001.

## Notes

### Competing Interest Statement

The authors have declared no competing interest.

